# LUCas: Light-Uncaged Cas13a using photocleavable interfering guide RNAs

**DOI:** 10.64898/2026.02.02.700737

**Authors:** Carlos F. Ng, Deepak Krishnamurthy, Andres Dextre, Aymeric Chorlay, Melanie Ott, Daniel A. Fletcher

## Abstract

CRISPR diagnostics have emerged as powerful tools for detecting infectious diseases, with the RNA endonuclease Cas13a enabling sensitive and specific, amplification-free RNA detection through collateral *trans*-cleavage of fluorescent reporters. However, background cleavage from unbound enzyme, contaminating nucleases, and unsynchronized initiation of reactions limits assay sensitivity and interpretability. A strategy to precisely control the onset of Cas13a catalytic activity, essentially a molecular “starting gun”, would address these challenges and expand assay design space. Here, we introduce Light-Uncaged Cas13a (LUCas), a light controllable system that directly gates Cas13a using a photocleavable interfering guide RNA (pc-igRNA) that suppresses *trans*-cleavage activity even in the presence of target RNA. Brief UV illumination releases this suppression, restoring full activity. Quantitative kinetic analysis reveals an approximately 100-fold suppression of *trans*-cleavage activity prior to photo-activation. Importantly, LUCas also suppresses target-independent background activity, enabling a predictive, background-limited determination of assay sensitivity. Using measured kinetic parameters, we predict and experimentally validate the limit-of-detection of the LUCas system. Finally, we demonstrate a multiplexed detection strategy termed “temporal barcoding,” which enables quantitative detection of viral co-infections in a single bulk reaction. Together, these results establish LUCas as a general framework for mechanistically informed, light-based control of Cas13a activity.

## 2 Introduction

Clustered Regularly Interspaced Short Palindromic Repeats (CRISPR)-based diagnostics have demonstrated great potential for sensitive and specific detection of DNA and RNA targets over the last decade. In particular, the CRISPR enzymes Cas12a and Cas13a have been used to develop a range of diagnostic platforms such as SHERLOCK, DETECTR, SATORI and CARMEN to detect nucleic acids from various pathogens and other targets [1, 2, 3, 4]. When Cas13a, a Class 2 type VI enzyme, complexes with a guide RNA (gRNA), it forms a ribonucleoprotein (RNP). The gRNA is a single-stranded RNA (ssRNA) containing a stem-loop structure important for enzyme recognition and a seed region complementary to a target of interest. Subsequent binding of target RNA to the seed region of the RNP results in a conformational change in Cas13a that unleashes non-specific cleavage of ssRNA, which is known as *trans*-cleavage activity. This *trans*-cleavage activity is mediated by two conserved HEPN (higher eukaryotes and prokaryotes nucleotide-binding) domains located on the solvent-accessible surface of Cas13a [1, 2, 5]. For diagnostic applications, the *trans*- cleavage activity is leveraged by adding a reporter molecule that consists of a fluorophore-quencher pair connected through a short ssRNA sequence, to the reaction. Upon target recognition by the gRNA, the RNP cleaves surrounding reporter molecules resulting in an increase of fluorescence signal over time, at a rate proportional to the target concentration [6].

To correctly interpret the signals generated in Cas13a in diagnostic assays, it is necessary to account for the various sources of noise that affects its signal-to-background ratio (SBR). For example, there is a background rate of cleavage of the reporter molecules even in the absence of a specific target RNA that can limit assay sensitivity. Cas13a reactions start as soon as a target RNA is bound, meaning that signal is being generated as soon as reagents are mixed together rather than when all target RNA has bound a Cas13a RNP and data collection is ready to begin. In addition, the uncontrolled nature of the amplification makes it challenging to distinguish the target-generated signal from that due to background nuclease activity. Lastly, when combined with pre-amplification techniques, the *trans*-cleavage activity of Cas13a can degrade intermediate products and hamper the detection sensitivity, thus requiring precise fluidic operations to mix the reaction at specific times [7, 8, 9].

Exploiting the full potential of Cas13a and other CRISPR/Cas enzymes for clinical applications requires strategies that can precisely control its activity in both space and time. In particular, a “starting-gun” for the *trans*-cleavage activity would be of practical value in designing and implementing diagnostic assays. To implement such a triggered initiation of the reaction, light-based control strategies have several key advantages: they are simple to implement, require no addition of reagents, and are highly tunable both in time and space.

Recently, there has been work demonstrating photo-control for Cas12a, motivated by the need to combine nucleic acid amplification methods with Cas12a-based detection in a single, one-pot assay without any fluid transfer steps [7, 10, 11, 12, 13]. In that case, activating Cas12a reactions at a specific-defined time prevents the *cis* and *trans*-cleavage activity from degrading the primers and products of the isothermal nucleic acid amplification steps, which would negatively impact the assay sensitivity [11]. The photo-control strategies developed for Cas12a involve the use of photocleavable ‘protective’ or ‘silent’ oligos that are complimentary to the gRNA hairpin structure [14], spacer region [15] or in both regions [7, 11, 16]. This prevents RNP formation and/or target binding, stalling the Cas12a reaction until light exposure occurs. Other approaches for controlling enzyme activity focus on the gRNA, which can be a more universal strategy. For example, some studies have demonstrated light-based control by embedding photocaged groups, creating a ‘cloaking’ effect that prevents target binding [10, 17, 18]. Other studies have extended the 3’ end of the gRNA for Cas12a with a steric hindrance effector [19] or an inhibitory extension [12]. Recently, allosteric control of Cas13d catalytic activity using light was demonstrated [20] and a photoactivatable split Cas13 for RNA base editing [21]. However, currently, a simple, light-based method for Cas13a that directly modulates enzyme *trans*-cleavage and is independent of RNP formation and guide-target kinetics is still lacking.

Here we present LUCas (Light-Uncaged Cas13a), a direct and modular photo-control strategy for Cas13a based on photocleavable interfering guide RNAs (pc-igRNAs). A pc-igRNA consists of a standard gRNA appended at its 5^*′*^ end with a single photocleavable linker followed by a short DNA interfering segment that suppresses *trans*-cleavage even in the presence of target RNA. Importantly, this design decouples control of enzyme activity from RNP assembly and guide–target binding. Through quantitative kinetic analysis, we show that Cas13a with pc-igRNAs operates in a well-defined suppressed state prior to illumination and transitions to a fully active state upon light exposure. We introduce a phenomenological “suppression factor” that captures the fold-change in *trans*-cleavage activity upon photo-activation. We demonstrate that this factor can be independently tuned by the length of the inhibitory DNA segment and the applied light dose. Notably, background cleavage is also suppressed before activation, enabling both predictive and experimental determination of the assay limit-of-detection, which we find to be set by intrinsic enzyme background activity.

To probe the mechanism underlying pc-igRNA mediated control, we use fluorescence anisotropy to track complex formation of the enzyme with pc-igRNAs and determine how photo-activation rapidly restores full Cas13a *trans*-cleavage activity. Finally, we leverage light as a temporal control knob to introduce “temporal barcoding,” a multiplexed Cas13a detection strategy in which distinct targets are encoded by their activation times rather than by spatial or spectral separation.

## 3 Methods

### 3.1 Protein purification

Protein purification was performed as previously described [22]. The LbuCas13a expression vector contains the codon-optimized Cas13a genomic sequence, N-terminal His6-MBP-TEV cleavage site sequence, and a T7 promoter binding sequence (Addgene Plasmid #83482). The protein is expressed in Rosetta 2 (DE3) pLysS E. coli cells in Terrific broth at 16 ^◦^C overnight. Soluble His6-MBP-TEV-Cas13a was isolated over metal ion affinity chromatography and the His6-MBP tag was cleaved with TEV protease at 4 ^◦^C overnight. Cleaved Cas13a was loaded onto a HiTrap SP column (GE Healthcare) and eluted over a linear KCl (0.25-1.0M) gradient. Cas13a-containing fractions were further purified via size-exclusion chromatography on a S200 column (GE Healthcare) in gel filtration buffer (20 mM HEPES-K pH 7.0, 200 mM KCl, 10% glycerol, 1 mM TCEP) and subsequently flash frozen for storage at −80 ^◦^C.

### 3.2 Guide RNA design

All gRNAs and pc-igRNAs were ordered from Synthego and Integrated DNA Technologies at a scale of 4 nmole (Table S1). The sequence was designed to recognize the SARS-CoV-2 N gene as previously described [22, 23].

### 3.3 Bulk Cas13a assay

Reactions were prepared by preassembling LbuCas13a-crRNA RNP complexes at 133 nM equimolar concentrations for 5 minutes at room temperature. After that, LbuCas13a is diluted to 25 nM in cleavage buffer (20 mM HEPES-Na pH 7.2, 50 mM KCl, 5 mM MgCl_2_ and 5% glycerol) with the addition of 400 nM of a polyU reporter molecule, 1 unit/µL Murine RNase Inhibitor (NEB, Cat#M0314), diethylpyrocarbonate (DEPC)-treated water (Fisher Scientific, Cat#BP561-1) and varying amounts of target concentrations. Once the reactions were mixed, the reactions were loaded into a 384-well (Corning, Cat#3544) and incubated in a plate reader (TECAN, Spark) for 60 minutes at 37 ^◦^C with fluorescence intensity measurements obtained every 2.5 minutes (*λ*_*ex*_: 485 nm; *λ*_*em*_: 535 nm). A calibration curve (Figure S6) was used to convert the relative fluorescence unit (RFU) to cleaved reporter concentration (in nM) as previously done [24]. The reaction rate (in nM/sec) is obtained by fitting a linear regression or the Michaelis-Menten equation to the cleaved reporter concentration from the initial time to the reaction duration time. For reactions using photocleavable guides, the “pre-stim” data was measured during the first 30 minutes of the reaction followed by the “post-stim” data measured for 1 hour.

#### UV stimulation apparatus

For UV stimulation, a 365 nm high-power 96-well LED array (Analytical, Lumidox II) was used to photoactivate individual 20-µL CRISPR reactions (Figure S3A). To apply a high UV dose in a 384-well plate, a custom acrylic plate adapter was used to align each of the 96 LEDs to be on top of individual wells (Figure S3B and Figure S3C). The 96-well LED array was connected to a controller unit that sets the power and stimulation time. A complete characterization of the illumination wavelengths and power level applied to an individual well can be found in Figure S4A and Figure S4B, respectively. The non-stimulated reactions were simply covered during activation by the custom adapter, which is designed to avoid premature activation in neighboring wells (Figure S5B). All measurements were conducted in triplicates.

#### Kinetic measurements

Bulk Cas13a cleavage kinetics were measured by adding RNA targets at defined concentrations to reaction mixtures containing Cas13a RNPs and fluorescent RNA reporters. Fluorescence intensity arising from reporter cleavage was monitored continuously for 30 min to quantify pre-stimulation activity. Reactions were then initiated by a 60 s exposure to 365 nm UV light using a Lumidox plate illuminator, followed by continued fluorescence measurements for an additional 60 min (post-stimulation phase).

Fluorescence time courses were converted to cleaved reporter concentration using calibration curves and analyzed using a bi-phasic Michaelis–Menten model that independently accounts for pre- and post-stimulation reaction regimes. Reaction rates were estimated from the fits to pre- and post-stimulation phases by extracting the time-scale (*τ*) predicted by the model-predicted cleaved reporter concentration curves.

Uncertainty in the estimated reaction rates was computed by combining two independent contributions: (i) variability across experimental replicates and (ii) uncertainty in the fitted model parameters. These contributions were combined in quadrature to yield the total uncertainty reported for each reaction rate.

#### Background activity measurements

To quantify target-independent background activity, reactions were performed at increasing RNP concentrations ranging from 25 to 250 nM. To reliably measure reporter cleavage at the low expected background catalytic rates, a higher reporter concentration (2000 nM) was used in these assays. Reaction rates were estimated using the same analysis pipeline as for the bulk kinetic measurements (Methods), by converting fluorescence signals to cleaved reporter concentrations using calibration curves and performing linear fits to extract reaction rates.

#### Fluorescence anisotropy assay

The photocleavable interfering guides (pc-igRNA) were labelled with Rhodamine Red C2 Maleimide (ThermoFisher, Cat#R6029) using the 5’ EndTag DNA/RNA Labeling Kit (Vector Laboratories, CA). After that, the fluorescently labelled pc-igRNA concentration were measured using a NanoDrop spectrophotometer (ThermoFisher) and aliquoated. For the actual measurements, a CRISPR-Cas13 reaction was prepared by mixing LbuCas13a with a 5’ fluorescently-labelled pc-igRNA using a ratio of 4:1. After a 5 minute incubation at room temperature, the mixture was diluted to a final concentration of 25 nM of LbuCas13a assembled with a photocleavable guide in cleavage buffer with 100 nM target RNA. Reactions were loaded into a 384-well plate (Corning, Cat#3820) and incubated in a temperature-controlled plate reader (TECAN, Spark) for 60 minutes at 37 ^◦^C with fluorescence polarization measurements taken every 2.5 minutes (*λ*_*ex*_: 570 nm; *λ*_*em*_: 635 nm) that were used to calculate anisotropy (r):

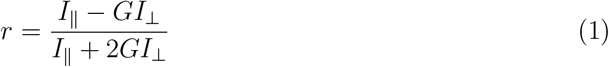

where *I*_∥_ and *I*_⊥_ are the background-subtracted parallel and perpendicular intensities, and *G* is a correction factor obtained from measuring polarization for a solution of 25 nM free floating Rhodamine in cleavage buffer. All measurements were conducted in triplicates.

For the simultaneous fluorescence and anisotropy measurements, the reactions were prepared as described previously and the plate reader perform an alternating sequence of fluorescence polarization and intensity measurements which were the pre-stimulation measurements. For the single stimulation using the plate reader, four consecutive fluorescence intensity measurements (*λ*_*ex*_: 360 nm; *λ*_*em*_: 520 nm) were used followed by a series of alternating fluorescence polarization and intensity measurements. For the pulsed stimulation, a single fluorescence intensity measurement was added after each alternating fluorescence polarization and intensity measurements. The instantaneous slope for each condition was calculated by measuring the slope between two consecutive points of the fluorescence intensity over their time difference.

## 4 Results

### 4.1 Design and demonstration of the LUCas System

In order to achieve light-based control of Cas13a activity, we designed a chimeric DNA-RNA photocleavable interfering guide (pc-igRNA). We previously introduced interfering guide RNAs (igRNA) that modulate the *trans*-cleavage activity of Cas13a through the addition of a 5’ DNA segment [25]. The photocleavable design consists of an interfering DNA segment linked to a canonical crRNA at the 5’ end via a photocleavable linker (Fig. 1A, Figure S1A). Since a sequence of repeating thymine (T) nucleotide bases is most effective at suppressing *trans*-cleavage activity, we designed and tested interfering DNA segments consisting of a 19T sequence. Upon RNP complex formation with Cas13a from *Leptotrichia buccalis* (LbuCas13a) in the presence of target RNA, the DNA segment of the pc-igRNAs interferes with the *trans*-cleavage activity leading to a “suppressed” state (Fig. 1A). When stimulated with a ultraviolet (UV) light dose at 365 nm, the resulting photocleavage releases the interfering DNA strand from the HEPN catalytic domain leading to a “photo-active” state (Fig. 1A).

**Fig. 1.**
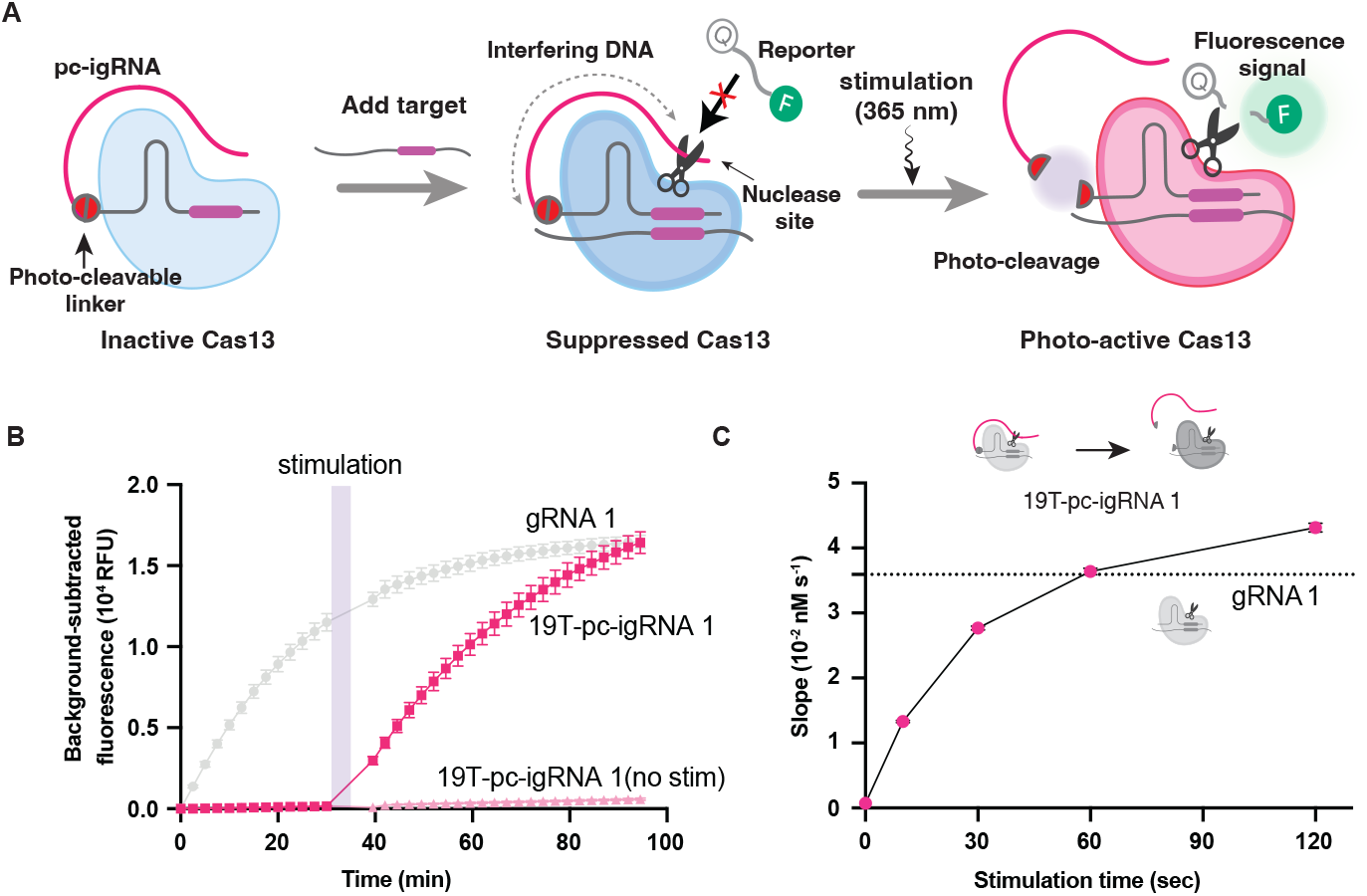
Demonstration of the LUCas system. **(A)** Schematic of the LUCas system for caging and light-triggered uncaging of Cas13a using a 19T photocleavable interfering guide RNA (19T-pc-igRNA). The pc-igRNA consists of a guide RNA (gRNA) linked at its 5^*′*^ end to a single-stranded DNA interfering segment via a photocleavable linker. **(B)** Photo-activation of Cas13a using a pc-igRNA containing a 19T interfering segment. Fluorescence increases due to reporter cleavage upon brief UV illumination (365 nm) in the presence of target RNA, whereas no UV exposure results in suppressed activity. Data points represent the mean across experimental replicates; error bars indicate standard error of the mean (SEM) (n=3). **(C)** UV dose–dependent activation of Cas13a, demonstrating that the extent of photo-uncaging quantitatively and precisely tunes the effective reaction rate. Data points and error bars represent the mean ± SEM for n = 3 wells.

We directly demonstrated both the suppressed and photo-activated states of the LUCas system by monitoring bulk fluorescence arising from cleavage of a fluorophore–quencher RNA reporter (5 uracils) [22] (Fig. 1B). In the presence of 10^6^ cp/*µ*L target RNA, a rapid increase in fluorescence was observed only in reactions receiving a 365 nm light dose, indicating light-triggered activation of Cas13a with pc-igRNA. In contrast, non-exposed reactions showed no measurable change in fluorescence, demonstrating strong suppression of *trans*-cleavage activity (Fig. 1B). As expected, reactions programmed with standard guides exhibited high cleavage activity prior to UV exposure and were unaffected by UV illumination (Fig. 1B). Importantly, we established the generality of the LUCas strategy by reproducing robust suppression followed by photo-activation across multiple guide and target RNA combinations (Figure S2).

Importantly, using light provides a tunable control knob for Cas13a activity in the LUCas system, as both illumination intensity and duration can be precisely modulated to control *trans*-cleavage. We find that increasing the 365 nm light dose photo-activates a larger fraction of the RNP population, producing a monotonic increase in reaction rate (Fig. 1C). This response saturates at approximately 60 s of UV exposure at an intensity of 16.1 mW*/*cm^2^ (total dose 96 mJ), beyond which no further increase in activity is observed and the reaction rate is comparable to that of a standard guide (Fig. 1C).

#### Characterization of enzyme *trans*-cleavage kinetics

As demonstrated in the previous section, the LUCas system allows the *trans*-cleavage activity of Cas13a enzyme to be rapidly switched from a suppressed to a photo-activated state using a brief light pulse. We next characterized the enzyme kinetics in these suppressed and photo-active states, a pre-requisite for exploring the design space of LUCas guides and enabling quantitative assays. We first conducted a sweep across 5 orders-of-magnitude in target concentration and measured the dynamics of cleaved reporter concentration during a pre-stimulation phase followed by a post-stimulation phase which was triggered by a 60 seconds UV light exposure (UV intensity: 16.1 mW*/*cm^2^, corresponding to full photo-activation, see Fig. 1C). For regular guide-target activation, the *trans*-cleavage activity has been shown to follow a Michaelis-Menten enzyme kinetics model [24]. Here, for the LUCas system, we hypothesized that the cleaved reporter dynamics could be predicted by a bi-phasic Michaelis-Menten enzyme kinetics model, with distinct kinetic parameters for the pre-and post-stimulation phases. This is given by the following equations for the pre- and post-stimulation phases, respectively (Fig. 2A):

**Fig. 2.**
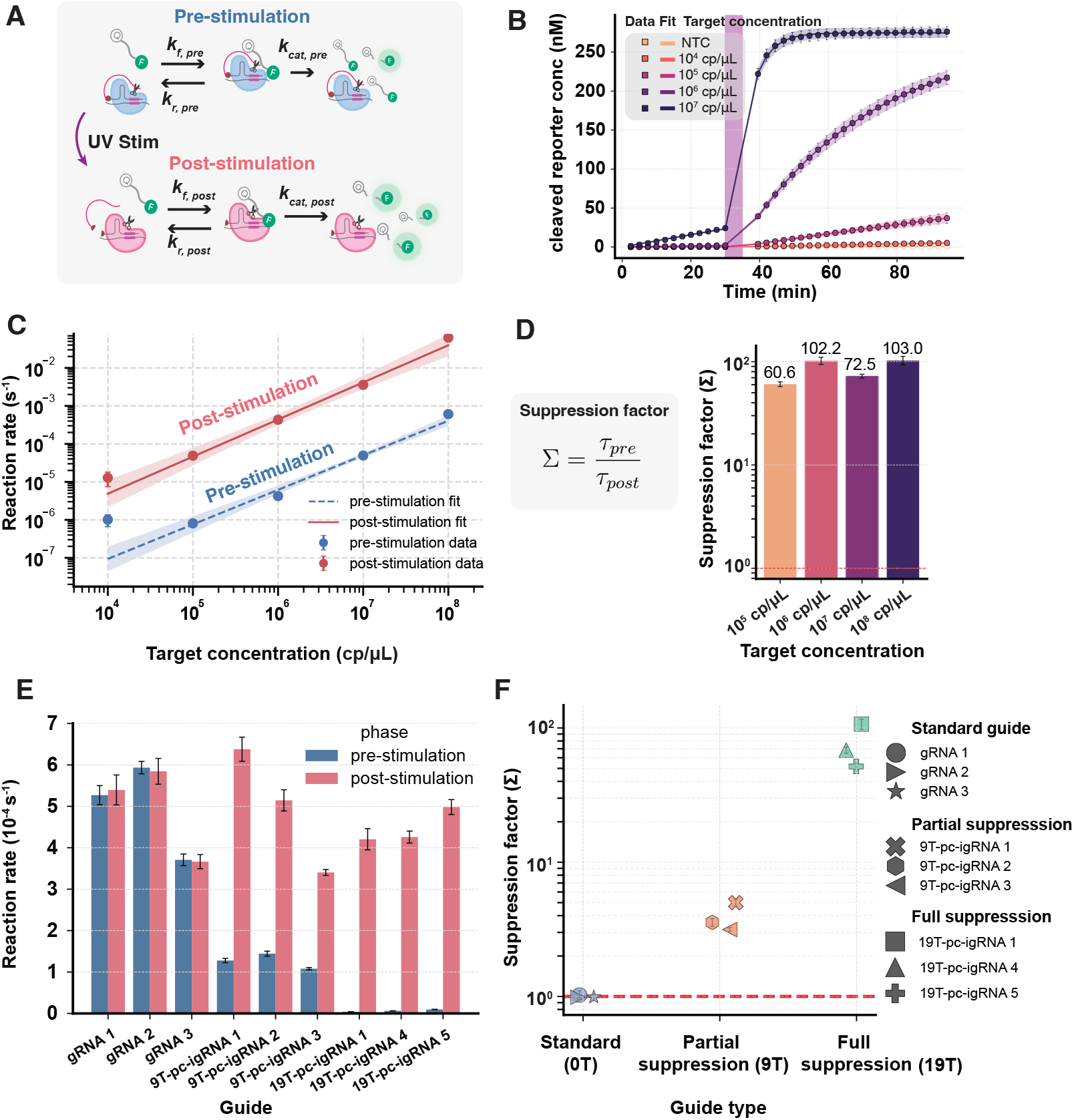
Kinetics of LUCas and Suppression Factor Measurements. **(A)** Schematic of *Lbu*Cas13a trans-cleavage kinetics using LUCas guides. **(B)** Cleaved reporter concentration as a function of time for increasing target concentrations using the 19T-pc-igRNA 1. Symbols represent experimental measurements (mean across replicates), and solid lines indicate fits to a two-phase Michaelis–Menten kinetics model. Shaded regions denote the 95% confidence interval (CI) of the fitted model. **(C)** Reaction rates extracted from the two-phase fits for the pre- and post-stimulation phases. Lines represent weighted linear regressions of the form Rate = (*k*_cat_*/K*_*M*_)[*E*], where [*E*] is the target concentration and *k*_cat_*/K*_*M*_ is the apparent catalytic efficiency. Shaded regions indicate the 95% CI of the fitted regression. **(D)** Suppression factor, defined as the ratio of post-to pre-stimulation reaction rates, plotted as a function of target concentration. **(E)** Pre- and post-stimulation reaction rates for guides containing increasing lengths of the interfering DNA segment (0T, 9T, 19T). **(F)** Corresponding suppression factors for guides with increasing interfering segment length. For panels **(B–F)**, error bars represent one standard deviation across independent experimental replicates.

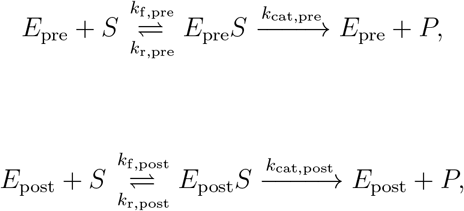

where *E* is the target-activated RNP complex, *S* is the substrate of uncleaved reporter molecules, *ES* is the RNP–reporter complex, and *P* is the cleaved reporter whose fluorescence we measure. The kinetic parameters *k*_f_ and *k*_r_ are the forward and reverse rates of *ES* formation, and *k*_cat_ is the catalytic turnover rate. Subscripts “pre” and “post” denote the parameters during the pre- and post-stimulation phases. ß

Under the assumptions of [*S*] ≪ *K*_*M*_, where *K*_*M*_ = (*k*_*r*_ + *k*_cat_)*/k*_*f*_ is the Michaelis– Menten constant, one can simplify the kinetics and write a closed-form solution for the cleaved reporter concentration. This assumption holds in our system since [*S*] ≈ 400 nM and *K*_*M*_ ∼ 𝒪 (100 nM to 1 *µ*M) (from previous measurements, see for instance [24]). This gives:

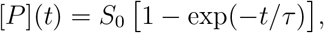

where *S*_0_ is the initial uncleaved reporter concentration and *τ* = *K*_*M*_ */*(*k*_cat_*E*_0_) is the characteristic reaction time scale. Here *S*_0_ is the initial reporter concentration and *E*_0_ the initial concentration of target-activated RNP.

Applying this model to the pre-stimulation phase one obtains:

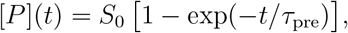

where *τ*_pre_ = *K*_M,pre_ */*(*k*_cat,pre_*E*_0_).

For the post-stimulation phase, which after *t* = *t*_stim_, we write:

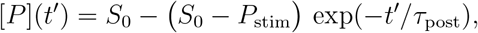

where *t*^*′*^ = *t* − *t*_stim_, *τ*_post_ = *K*_M,post_*/*(*k*_cat,post_*E*_0_), and *P*_stim_ is the cleaved reporter concentration at the stimulation time *t*_stim_.

We performed non-linear least-squares fitting of the pre- and post-stimulation data and observed excellent agreement with this biphasic Michaelis–Menten model (Fig. 2B). From the fits, we extracted the characteristic reaction time scales *τ*_pre_ and *τ*_post_ as functions of the target concentration (Fig. 2C). Over 4 decades in target concentration, both pre- and post-stimulation reaction time scales exhibited a linear dependence on target concentration (*R*^2^ = 0.99 for pre-stimulation and *R*^2^ = 0.95 for post-stimulation)(Fig. 2C). Thus, as predicted by the Michaelis-Menten model, the corresponding reaction rates satisfy 1*/τ*_pre_ = (*k*_cat,pre_*/K*_M,pre_) *E*_0_ and 1*/τ*_post_ = (*k*_cat,post_*/K*_M,post_) *E*_0_.

From linear fits of reaction rate versus target concentration, we estimate

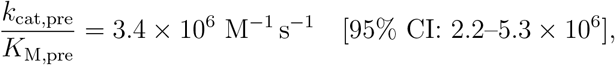

and

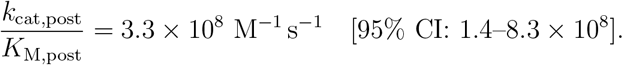

We define the suppression factor, Σ, as the ratio of post-to pre-stimulation enzyme catalytic efficiency, Σ = (*k*_cat_*/K*_*M*_)_post_*/*(*k*_cat_*/K*_*M*_)_pre_, or equivalently as the ratio of the characteristic reaction time scales, Σ = *τ*_pre_*/τ*_post_. For the 19T-pc-igRNA, we measure Σ ≈ 10^2^, corresponding to an approximately hundred-fold increase in activity upon illumination (Fig. 2D). Notably, Σ is largely independent of target RNA concentration (Fig. 2D), indicating that suppression and photo-activation are governed by guide caging rather than guide–target binding kinetics.

To confirm this independence, we next measured the pre-and post-stimulation reactions rates across a range of different guide-target combinations and for two different lengths of the interfering DNA segment, one consisting of 9 and, the other consisting of 19 thymine (T) bases. We label these two types of interfering guides as 9T-pc-igRNA and 19T-pc-igRNA corresponding to partial and full interfering activity from the DNA segment. As expected, canonical guides with no interfering DNA segment or photocleavable linker showed the same reaction rates pre-and post-stimulation, and suppression factors Σ ≈ 1 (Fig. 2E). Partially interfering guides (9T-pc-igRNA) allowed the enzyme to retain some activity pre-stimulation and then, gain its full activity upon photo-activation, while fully interfering guides (19T-pc-igRNA) showed the highest suppression of activity prior to photo-activation (Fig. 2E). Importantly, while absolute reaction rates varied across different guide–target combinations, the measured suppression factors clustered into logarithmically separated regimes. These correspond to Σ ≈ 1 (no suppression for standard guides), Σ ≈ 3–5 (partial suppression for 9T-pc-igRNAs), and Σ ≈ 50–125 (full suppression for 19T-pc-igRNAs) (Fig. 2F). These results demonstrate that the photo-activation and associated suppression factors are a strong function of the interfering DNA length and are relatively insensitive to the guide-target combination, demonstrating a direct light-based control of the *trans*-cleavage activity of the enzyme.

#### Suppression of non-specific enzyme activity and limit-of-detection measurements using LUCas

In the previous section, we demonstrated that the suppression and photo-activation of LUCas guides, in the presence of target RNA, could be characterized via a suppression factor based on the kinetic rates. We next characterized the impact of suppression and photo-activation on the non-target-dependent background activity of the enzyme. This analysis leads to and culminates in our ability to predict and measure the limit-of-detection of a Cas13a reaction using a single guide with the LUCas system.

To characterize background rates, we measured the dynamics of the cleaved reporter concentration over a range of RNP concentrations. For standard guides, we observed a steady linear increase in cleaved reporter concentration with time, at a rate that is RNP dependent and clearly greater than the no-RNP control (Fig. 3A). This no-RNP rate represents the noise floor of our measurements and results from non-enzyme-related background processes that may include reporter degradation [26]. In contrast, for the 19T-pc-igRNA we detected suppressed background activity in the pre-stimulation phase, followed by an increased rate after photo-activation (Fig. 3B). Importantly, this rate change after stimulation occurred only when RNP was present, which rules out photo-induced increase in reporter degradation as a possible reason for this change. Reaction rates extracted from least-squares fits to the cleaved reporter concentration data showed linear scaling with RNP concentration for both pre- and post-stimulation phases, with the post-stimulation background rate being comparable to the standard guide case, as expected (Fig. 3C). We can therefore model the RNP-dependent background reaction rate as:

**Fig. 3.**
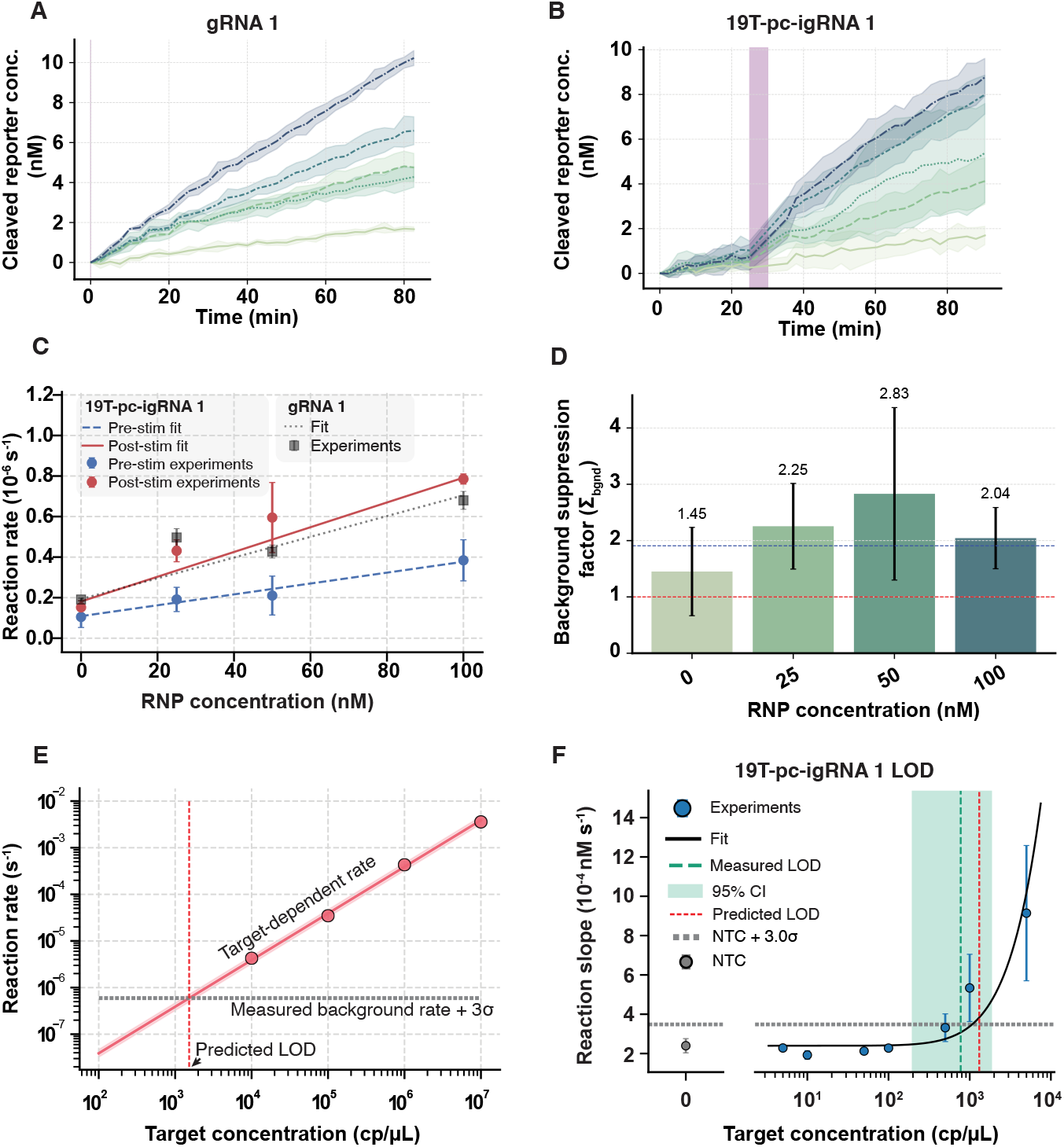
Background kinetics of LUCas and limit-of-detection measurements. **(A,B)** Cleaved reporter concentration as a function of time showing background trans-cleavage activity of the RNP with standard guides and pc-igRNAs, respectively, in the absence of target RNA. Solid lines indicate the mean across experimental replicates, and shaded regions denote the 95% confidence interval (CI). **(C)** Reaction rates extracted from linear fits to the pre- and post-stimulation phases of the curves shown in **(A,B)**, enabling quantification of background activity. **(D)** Background suppression factor, defined as the ratio of post-to pre-stimulation reaction rates in the absence of target RNA, demonstrating that LUCas guides suppress background activity by approximately two-fold. **(E)** Post-stimulation reaction rate as a function of target concentration (red symbols with linear fit), shown together with the detection threshold derived from background rates measured at 25 nM RNP (mean + 3*×* SD; grey dashed line). Mapping this threshold onto the fit yields the predicted limit-of-detection (LOD) (vertical red dashed line). **(F)** Experimental LOD measurements for the 19T-pc-igRNA 1 guide. Post-stimulation reaction rates were fitted with a three-parameter logistic model (solid black curve). The experimentally determined LOD is indicated by the median estimate (vertical green dashed line) with its 95% CI (green shaded band), shown alongside the predicted LOD (vertical red dashed line). For panels **(C,D,F)**, error bars represent the standard error, obtained by combining variability across experimental replicates and uncertainty from the fitted rates in quadrature.

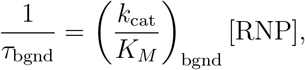

where the background catalytic efficiencies are found to be

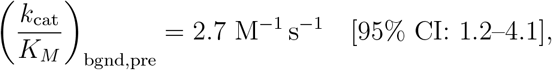

and

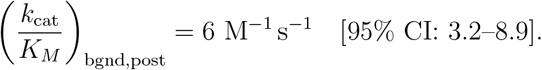

Thus, the LUCas guides suppress Cas13a RNP background activity by roughly two-fold, again in a manner that is independent of RNP concentration (Fig. 3D).

Using these measurements of background activity, we next established the limit-of-detection (LOD) of the LUCas system. Since our analysis is rate-based, we defined the detection threshold as

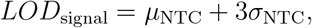

where *µ*_NTC_ and *σ*_NTC_ are the mean and standard deviation of the reaction rate for the non-target control (NTC), computed across three replicate wells. The LOD in signal space is reached when the target-dependent rate equals this threshold, i.e. 1*/τ*_post_ = *LOD*_signal_. This implies that the LOD in concentration space is given by

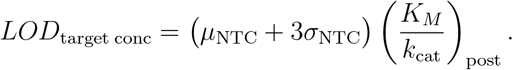

Using the above model and our measured target-dependent and background kinetic parameters, we predict a limit of detection (LOD) of 1500 copies*/µ*L (95% CI: 1250–1800) for a single reaction (Fig. 3E). We next determined the LOD experimentally by measuring post-stimulation reaction slopes across increasing target concentrations for the 19T-pc-igRNA 1. The slope data were fit using a three-parameter sigmoid model [27], and the same NTC-based LOD threshold was applied (Fig. 3F). Mapping the resulting threshold from reaction-rate space to concentration space yielded an experimental LOD of 815 copies*/µ*L (95% CI: 200– 1881). Notably, the predicted and experimentally measured LOD values are in agreement, as evidenced by their overlapping confidence intervals, indicating that the LOD of the current LUCas system is set by the background catalytic activity of the enzyme (Fig. 3F). This conclusion is further supported by measurements of two additional guide–target combinations using the same 19T interfering segment, both of which yielded comparable LOD values of 430 copies*/µ*L (95% CI: 250–900) and 520 copies*/µ*L (95% CI: 12–730) (Figure S7).

#### Mechanism of suppression and photo-activation in pc-igRNAs

The LUCas system leverages pc-igRNAs, which are hypothesized to work by blocking RNA cleavage through the *trans*-cleavage catalytic domain on Cas13a [25]. Molecular dynamics (MD) simulations predict that the ssDNA segment attached to the 5’ end of the guide is held close to the Cas13a surface in a manner that colocalizes the ssDNA segment with the catalytic HEPN domain (Fig. 4A). To test whether pc-igRNAs complex effectively with Cas13a and photocleaving releases the interfering DNA segment, we used fluorescence anisotropy measurements of the molecular rotational rate of the partially-blocking 9T-pc-igRNA to assess their binding to and release from Cas13a (Fig. 4B).

**Fig. 4.**
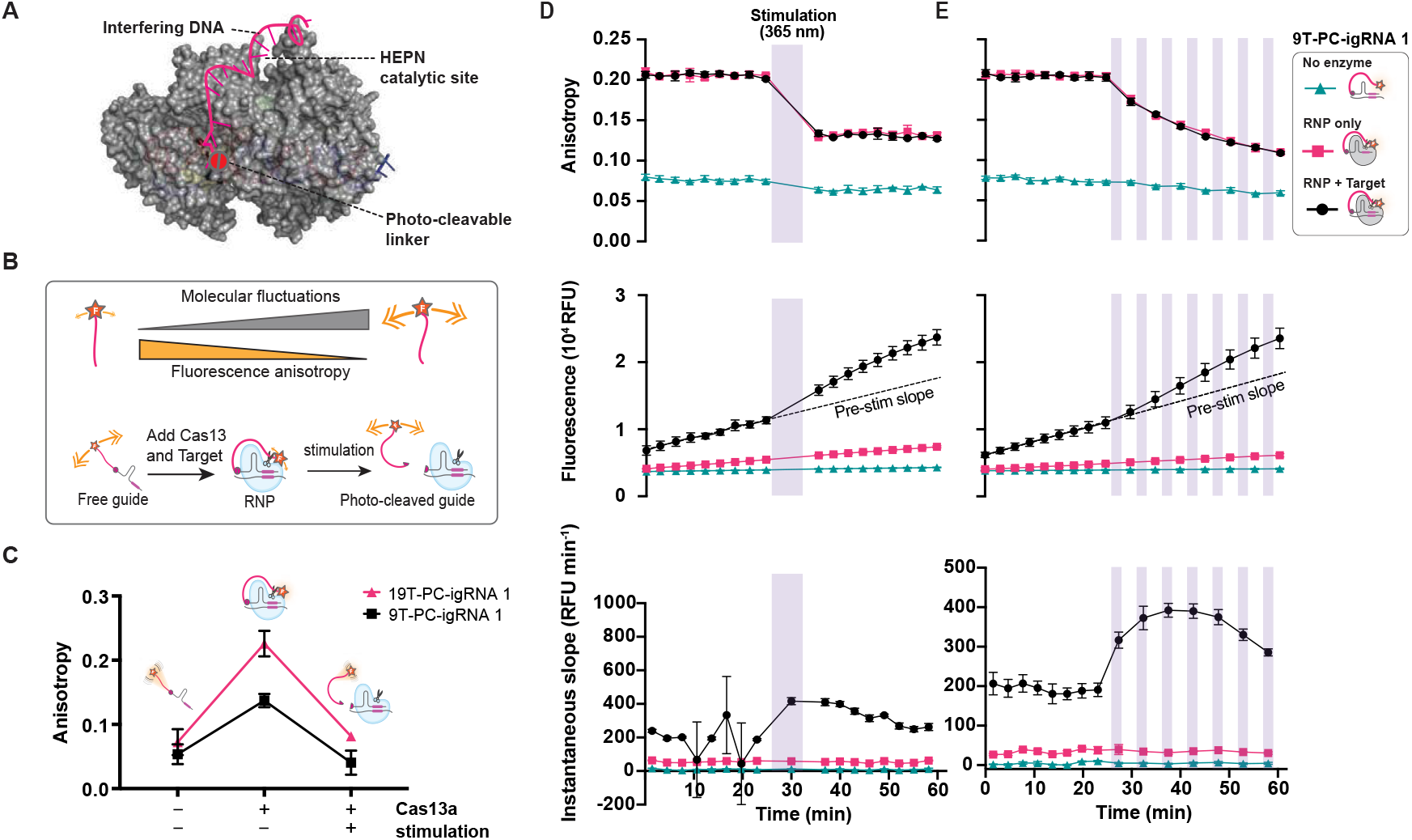
Mechanism of photo-activation using photocleavable interfering guide RNAs. **(A)** Structure of the LbuCas13a:19T-pc-igRNA:target RNA complex. **(B)** Schematic representation of the fluorescence anisotropy measurements used to elucidate the mechanism for LuCas. **(C)** Fluorescence anisotropy measurements using 5’ fluorescently-labelled gRNA for different lengths of the interfering DNA segment. Freely-suspended CRISPR guides (gRNAs) lead to a low fluorescence anisotropy value, while gRNAs complexed with LbuCas13a, which forms a ribonucleoprotein (RNP), leads to a higher anisotropy value. Upon UV illumination, the anisotropy relaxes to a lower value when compared to the RNP case. This observation is consistent with photo-release of the interfering ssDNA segment from the RNP complex. Data points show mean ± standard deviation (SD). **(D)** Simultaneous temporal measurements of fluorescence anisotropy and LbuCas13 *trans*-cleavage activity (via cleaved reporter intensity) shows the photo-activation of LbuCas13a by a single UV dosage (365 nm). The change in *trans*-cleavage rate occurs concurrently with the change in fluorescence anisotropy. **(E)** Simultaneous temporal measurements of fluorescence anisotropy and LbuCas13 *trans*-cleavage activity shows that the photo-activation of LbuCas13a can be precisely-tuned by pulsed UV illumination. Each UV pulse photo-activates a new population of LbuCas13a, as confirmed by a concurrent change in anisotropy. In panels (D)-(E) error-bars represent the standard error of the mean across three replicates.

Fluorescence anisotropy has been a powerful tool for quantifying protein binding, protein denaturation and protein dynamics, where it provides quantitative information about rotational degrees of freedom [28]. More recently, this technique has been used to study gRNA-target RNA binding affinities [29]. Small molecules such as gRNA that are free in solution have rapid rotational rates and hence low anisotropy values, while molecules bound to larger proteins have slower rotational rates leading to higher anisotropy values [28]. We measured the fluorescence anisotropy of a 5’ fluorescently-labeled 9T-pc-igRNA before and after UV stimulation, and we simultaneously measured the *trans*-cleavage activity to establish a mechanism for LUCas.

For the 19T-pc-igRNA without Cas13a, we measured low anisotropy values corresponding to the rotational rate of free molecules (Fig. 4C). Once Cas13a and the target RNA are added, we found a significant increase in anisotropy indicating a stable RNP complex formation (Fig. 4C). Using two versions of pc-igRNAs (one with a 9T and the other with a 19T interfering segment), we found that the increase in anisotropy is greater for the longer guide (19T) than the shorter one (230% for 19T vs 157% for 9T). Using the average size of 0.5 nm per nucleotide and the total number of nucleotides on the interfering strand, the total length is estimated to be 45 Å for the 9T and 95 Å for the 19T [30]. These data are consistent with a picture in which the longer DNA segment spans the distance between the maturation site and HEPN domains on Cas13a (distance ≈ 50 Å) and, hence, more effectively interferes with reporter binding and catalytic activity, consistent with increased suppression by these guides (Fig. 4A, Fig. 2E and Fig. 2F). Upon subsequent UV exposure, we found that the measured anisotropy fell back to values similar to those of free guide in solution, indicating efficient photocleavage and release of the interfering DNA segment from Cas13a (Fig. 4C).

How do these changes in anisotropy correlate with the *trans*-cleavage activity of Cas13a? To understand this we simultaneously measured the fluorescence anisotropy and *trans*-cleavage activity in the same experiment. We found that a single exposure of UV light (duration: 40 s, UV dose: 160 µJ) led to a step decrease in fluorescence anisotropy for 9T-pc-igRNA with Cas13a, independent of the presence of target RNA. In contrast, no change in anisotropy was observed for the case of free guides, even in the presence of target RNA (Fig. 4D). This step decrease in anisotropy temporally corresponded with a step increase in *trans*-cleavage activity (Fig. 4D).

When exposed to several shorter exposures of UV light (duration: 10 s, UV dose: 40 µJ), we observed rapid, step decreases in fluorescence anisotropy that corresponded with step-increases in *trans*-cleavage activity (Fig. 4E), consistent with additional populations of the RNP being photo-activated. The response eventually saturates due to the consumption of uncleaved reporter molecules in solution (Fig. 4E, bottom).

Our results from these fluorescence anisotropy measurements indicate that pc-igRNAs effectively form RNPs despite the presence of the interfering DNA segment at the 5’ end of the guide. Furthermore, we confirm that recovery of *trans*-cleavage activity from a suppressed state upon UV stimulation results from rapid photocleavage and subsequent release of the interfering DNA segment from the vicinity of the HEPN catalytic domain. This release process is faster than the time-scale of our measurements of both reporter cleavage and anisotropy, as we did not capture the transition from off to on. Our anisotropy measurements therefore highlight a key feature of the LUCas system: strong Cas13a binding and suppression by the pc-igRNAs followed by rapid release of the interfering DNA and restoration of activity after photocleavage.

#### Temporal barcoding with the LUCas system

Pathogen co-infections in a single host are common in nature and in clinical settings [31, 32, 33]. For that reason, multiplexing is a desired feature in most diagnostics assays. For CRISPR-based diagnostics, this is possible using different gRNA that target distinct pathogens or viruses. However, an outstanding issue is distinguishing the activity from different guides targeting different pathogens in a single bulk reaction, since modest sequence dependence of the *trans*-cleavage activity limits assays to using a single channel for reporter fluorescence.

In earlier work, we demonstrated one type of multiplexing strategy by leveraging the distinct kinetic signatures of different guides and interfering guide lengths, which we referred to as ‘kinetic barcoding’ [25]. Here, we leveraged the LUCas system to introduce an alternative multiplexing strategy that we call ‘temporal barcoding’. This strategy enables separately identifying the *trans*-cleavage activity of two or more different guides (e.g., one standard gRNA and one 19T-pc-igRNA) in a single CRISPR reaction using light to temporally control when a reaction is activated (Fig. 5A). Initially, in the simplest case of two different target RNAs of interest, only the RNP with the standard guide is active and the RNP with the 19T-pc-igRNA is suppressed. After a fixed duration of time, the reaction is exposed to UV light, activating the RNP with the 19T-pc-igRNA. This will result in a unique change in fluorescence signal over time, or a temporal barcode, that depends on the presence or absence of the two targets.

**Fig. 5.**
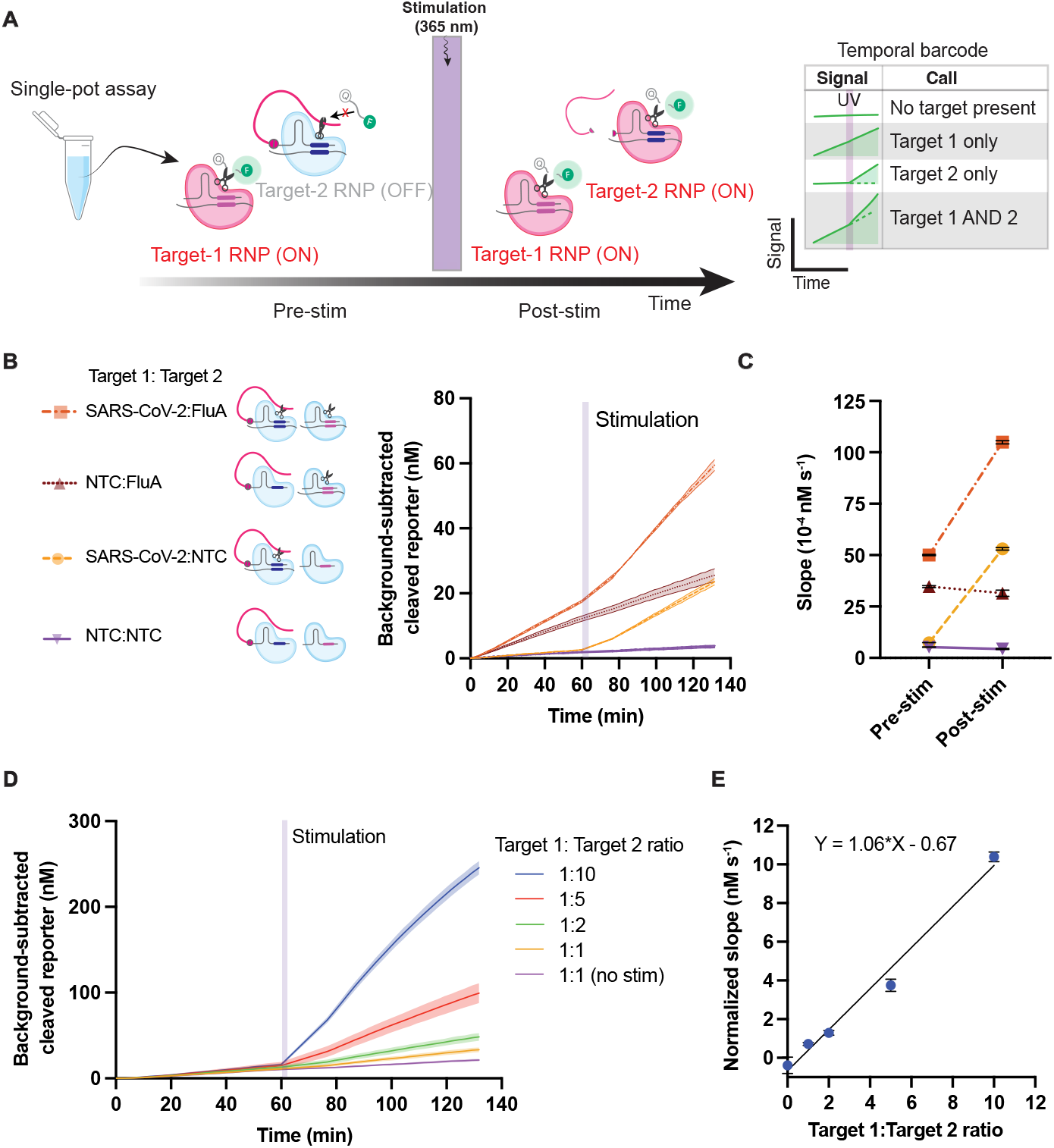
LUCas enables temporal barcoding assays for detecting multiple targets in a single reaction. **(A)** Schematic representation of the temporal barcoding assay, which enables differentiation between the *trans*-cleavage activity of one standard guide RNA (gRNA) and one photocleavable guide (pc-igRNA) in a single CRISPR reaction. **(B)** Demonstration of the LUCas assay for detecting two viruses (FluA and SARS-CoV-2) in a single reaction using a temporal-barcoding approach. The standard guide targets FluA (gRNA FluA), while the photocleavable guide is complementary to a segment of the SARS-CoV-2 virus genome. The curves are shown as a mean of three replicates and the shaded area around the curves show the SEM. **(C)** The distinct temporal barcodes, based on measured rate changes (pre-stim and post-stim), allow distinguishing between the four possible outcomes in a single reaction. Error bars represent the mean ± SEM (n=3). **(D)** Measured cleaved reporter concentration over time of a reaction with a mixture different ratios of target concentrations. Error bars represent the SEM (n=3). **(E)** Scatter plot of normalized slope difference 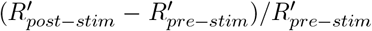, where *R*^*′*^ is the reaction slope measured using a linear fit, and ratio of target concentrations shows a linear relationship, as expected. This relationship can be used to quantitatively estimate the target concentration ratios of a sample, given knowledge about individual guide-dependent activity. Data points show the mean ± SEM (n=3).

To demonstrate this method, we combined a mixture of Influenza A (FluA) and SARS-CoV-2 whole genome RNA in one CRISPR reaction using a standard gRNA for FluA and a 19T-pc-igRNA for SARS-CoV-2. The measured fluorescence intensity and slope (before and after UV illumination) are shown in Fig. 5B and Fig. 5C, respectively. We find that reaction slopes pre and post-UV can be temporally segmented based on the presence of one or more targets (Fig. 5C). This demonstration shows how temporally separating the activity of two guides in a single reaction enables multiplexed detection of two targets using LUCas. To test the practically relevant case where the concentrations of the two targets are different, we varied the 19T-pc-igRNA target concentration and measured the cleaved reporter concentration (Fig. 5D) along with the pre-stim and post-stim reaction rates (Fig. 5E). The post-stimulation reaction slopes showed an increase that was dependent on target concentration ratio (Fig. 5E). In addition, the normalized slope difference 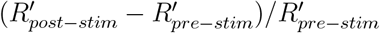, where *R*^*′*^ is the reaction slope, showed a linear relationship with the ratio of target concentrations, as expected (Fig. 5E). Furthermore, the slope of the linear fit was found to be proportional to the ratio of activities of the two guides for a given target concentration. This implies that one can quantitatively estimate the concentrations of at least two different targets in a single CRISPR reaction using this technique.

## 5 Discussion

In this work we introduced the LUCas system, which allows direct optical control of Cas13a *trans*-cleavage activity, independent of RNP formation and guide-target binding kinetics. LUCas allows modulation of the catalytic output of the enzyme in the suppressed and photo-activated states without perturbing the molecular recognition steps; in other words, suppression is present even when target RNA is bound to the RNP. Upon brief light exposure, the Cas13a activity rapidly rises to the full rate expected for a given guide-target combination.

We characterized the kinetics of the LUCas system and find that it follows a bi-phasic Michaelis-Menten model with distinct kinetic parameters for the pre- and post-stimulation phases. Importantly, this change in kinetic rates can be captured through a generalizable phenomenological suppression factor, which is the ratio of post-stimulation to pre-stimulation reaction rates, and it is independent of target concentration and guide-target combinations. We find that this suppression factor is 100-fold for the longest interfering DNA segment tested (19T) and can be tuned by two independent control knobs, namely the interfering DNA length and light dose, respectively. Furthermore, we find that the background activity of the enzyme in the absence of target RNA is also suppressed by a factor of 2, thus reducing the background above which a positive signal must rise to be detected. The bi-phasic kinetic model and the phenomenological suppression factor that we introduce could be a useful general framework to quantify how enzyme activity is modulated in other gating strategies, including alternative light-control methods.

Using our measurements of both target-dependent and background kinetic rates, we quantitatively predict the limit-of-detection of the LUCas system to be on the order of 1e3 cp/µL, and find excellent agreement with experiments across three different guide-target combinations. Lastly, using the light-controlled features of the LUCas system, we introduce a multiplexing strategy that we call ‘temporal barcoding’ and use it to detect a co-infection of SARS-CoV-2 and Influenza-A in a single bulk reaction, using light to sequentially activate two populations of RNPs, each one targeting a specific virus. Further, we showed that the relative abundance of two viral RNAs can be estimated in a single bulk reaction.

In contrast to earlier light-control strategies for CRISPR enzymes, the LUCas system directly gates enzymatic *trans*-cleavage activity, with or without the target bound, rather than perturbing RNP formation or suppressing target binding. This strategy avoids the high background activity of unbound or Apo-Cas13a, reduces background activity of the RNP, and allows time for all target RNA to bind to the RNP complex before the reaction starts. In terms of chemical modification, LUCas requires only a single photocleavable molecule per guide (compared to three or more in Cas12a light-activation strategies [7, 10, 11, 14]), and it does not perturb guide structure or guide–target binding kinetics, thereby simplifying assay design and alleviating complex optimization requirements. While we focused on amplification-free detection in this work, we expect LUCas to retain its key advantages when combined with amplification steps such as rolling circle amplification (RCA) and loop-mediated isothermal amplification (LAMP). These advantages include the ability to quench Cas13a *trans*-cleavage activity until the final detection step and to prevent degradation of amplification substrates and products, potentially leading to improved sensitivity.

While we have already demonstrated LUCas for the detection of two different targets, using temporal barcoding of distinct targets, the system is well-positioned for further multiplexing strategies. In particular, different targets can be encoded by including photocleavable moieties that are cleaved at distinct wavelengths [34]. LUCas can also be combined with other multiplexing strategies including a light-triggered form of “kinetic barcoding” [25], wherein the photocleavable molecule can be placed at different locations at the 5’ end of the gRNA to enable distinct and well-charactetrized suppression factors for different gRNA populations aimed at detecting different targets. LUCas also lends itself to spatial barcoding of different guides. This could be implemented on patterned surfaces or beads, with the LUCas system activating Cas13a *trans*-cleavage activity where and when needed. Finally, to improve implementation accessibility, modular design strategies such as click chemistry could be used to convert regular gRNAs into pc-igRNAs [35].

A defining feature of the LUCas system is the combination of strong enzymatic suppression with rapid activation, such that Cas13a binding and catalytic output can be cleanly decoupled. Further study of this behavior will help clarify how secondary interactions between the interfering DNA segment and surface residues of Cas13a modulate accessibility of the HEPN catalytic domains. In tandem, statistical mechanics and molecular dynamics models could provide a predictive framework for how interfering segment length, flexibility, and sequence set the degree of suppression and the kinetics of activation after photocleavage. Engineering Cas13a, and the pc-igRNAs to exploit these interactions more effectively has the potential to further enhance suppression, thereby improving signal-to-background ratios and expanding the performance envelope of Cas13a-based detection systems.

## 6 Acknowledgments

We thank all members of the Fletcher and Ott laboratories for feedback, discussions, and assistance. We also thank Professor Sungmin Son for providing us with some guide sequences that were used in this study.

## 7 Author contributions

C.F.N.: Conceptualization, Formal analysis, Investigation, Validation, Visualization, Writing – original draft. D.K.: Conceptualization, Data curation, Formal analysis, Methodology, Validation, Visualization, Writing – original draft. A.D.: Methodology, Visualization, Writing – review & editing. A.C.: Methodology, Writing – review & editing. M.O.: Funding acquisition, Resources, Supervision, Writing – review & editing. D.A.F.: Conceptualization, Funding acquisition, Resources, Supervision, Project administration, Writing – review & editing.

## 8 Supplementary data

Supplementary data is available online.

## 11 Data Availability

All data supporting the findings of this study, including raw fluorescence time-series, processed kinetic measurements, analysis scripts, and figure-generation code will be publicly available.

