## Supplemental Information for "LUCas: Light-Uncaged Cas13a using photocleavable interfering guide RNAs"

**Contents**

Figure S1: Photocleavable linker chemical structure.

Figure S2: Repeatability and reproducibility of LUCas guides.

Figure S3: LUCas assay illumination hardware.

Figure S4: Characterization of the Lumidox LED array.

Figure S5: No activity in non-stimulation wells.

Figure S6: Fluorescence versus cleaved and uncleaved reporter concentration calibration.

Figure S7: Limit-of-detection analysis for additional photo-cleavable guides.

Table S1: Oligonucleotides used in this study.

### Supplementary Figures

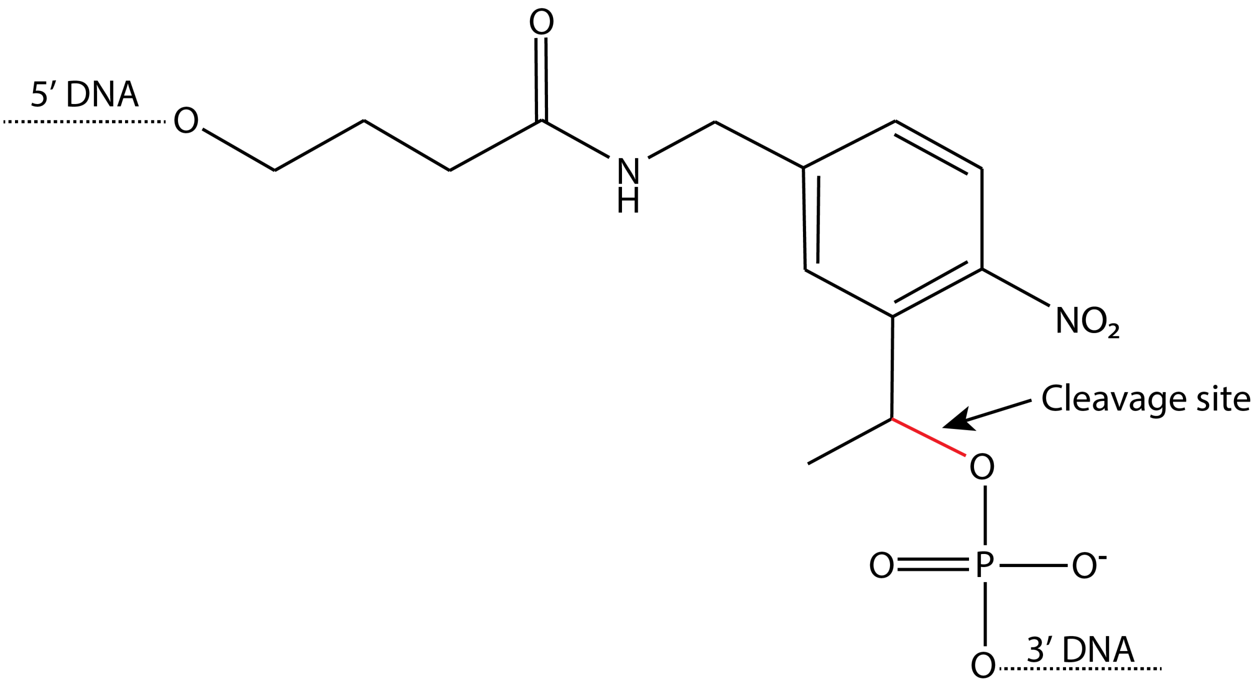


**Figure S1: Photocleavable linker chemical structure. (A)** Photocleavable Linker located in the DNA region of the interfering guide RNA (igRNA).

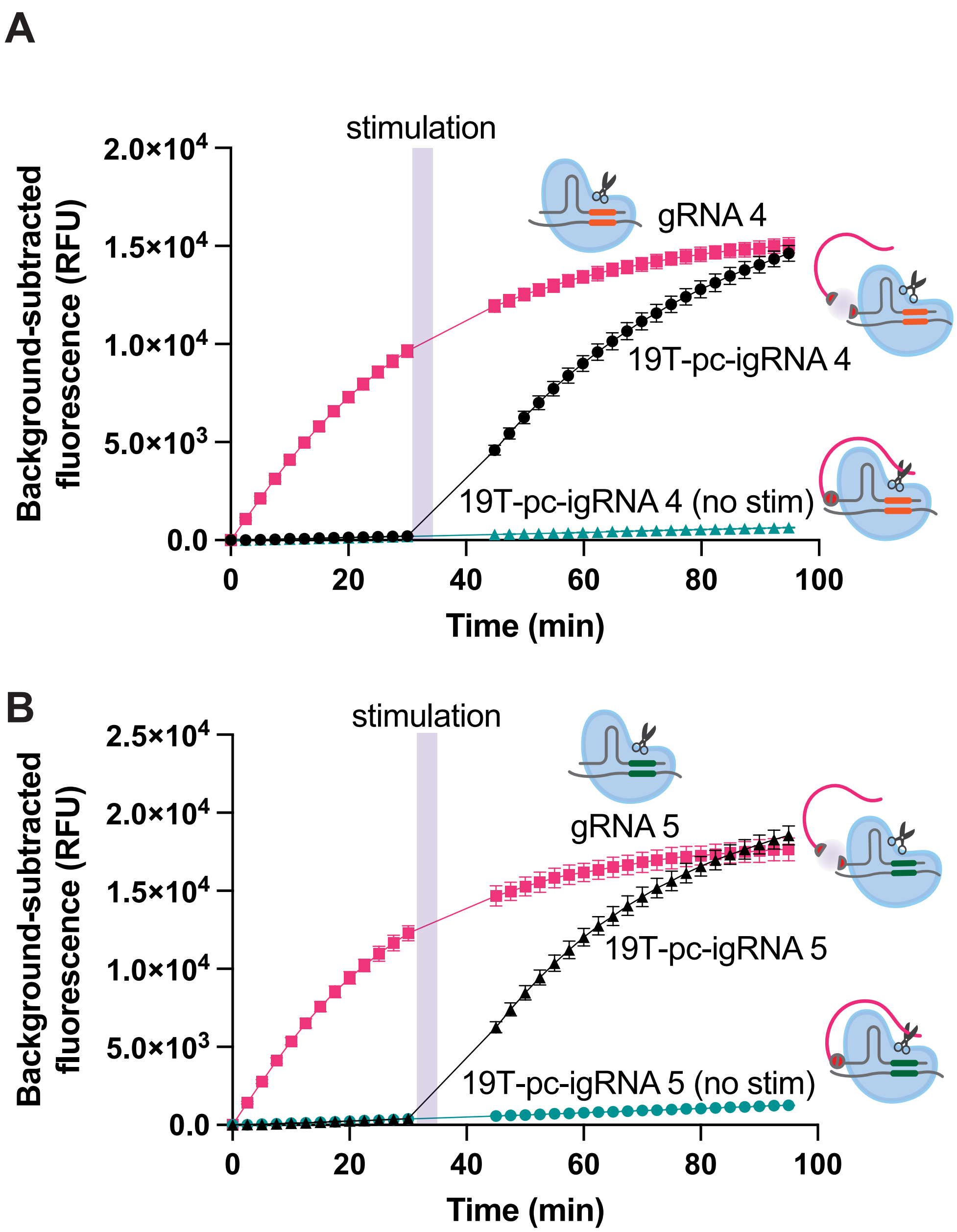


**Figure S2: Repeatability and reproducibility of LUCas guides.** Fluorescence measurements over time were performed for two guides with different spacer regions: **(A)** 19T-pc-igRNA 4 **(B)** 19T-pc-igRNA 5. Data points represent the mean ± standard error of the mean (SEM) (n=3). The same RNA target concentration (1e6 cp/µL) was added for each condition.

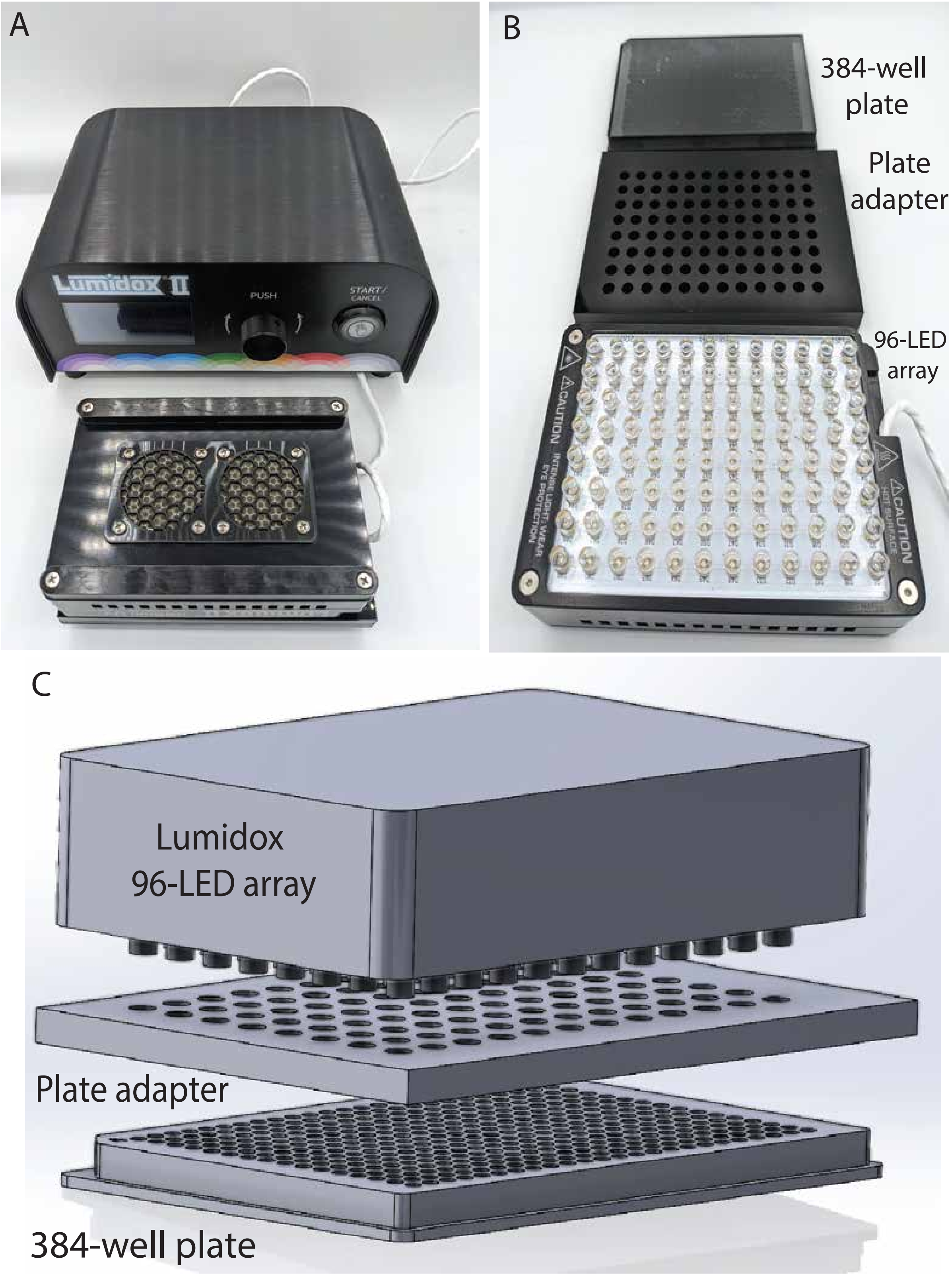


**Figure S3: LUCas assay illumination hardware. (A)** Lumidox II Controller connected to the 96-LED array (365 nm). **(B)** 96-LED array, acrylic custom plate adapter and a 384 well plate. **(C)** The 96-LED array is placed on top of an acrylic custom plate adapter which aligns the individual LEDs directly above a single well from a 384-well plate. All non-exposed wells are used as no stimulation wells.

A

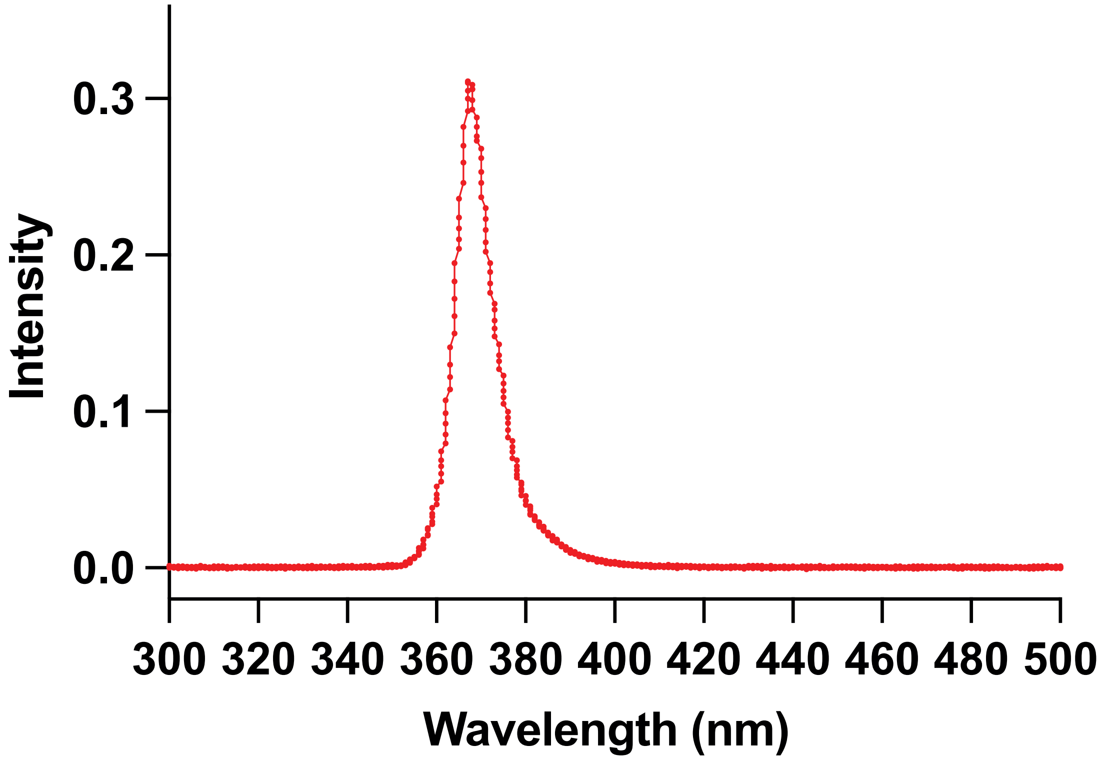


B

| Power setting on Lumidox (mW) | Irradiance per well (mW/cm$^2$) | Measured Power at exposed well (mW) | Measured Power at non-exposed well (mW) |
|---|---|---|---|
| 20 | 2.26 | 0.226 | 1.15e-3 |
| 40 | 5.70 | 0.570 | 2.23e-3 |
| 60 | 9.18 | 0.918 | 3.34e-3 |
| 80 | 12.70 | 1.270 | 4.46e-3 |
| 110 | 16.12 | 1.612 | 5.56e-3 |

**Figure S4:** **Characterization of the Lumidox LED array.** **(A)** Spectrum of a single LED from the Lumidox 96-LED Array. **(B)** Summary of measured power for both exposed and non-exposed single wells in a 384 well plate.

**A**

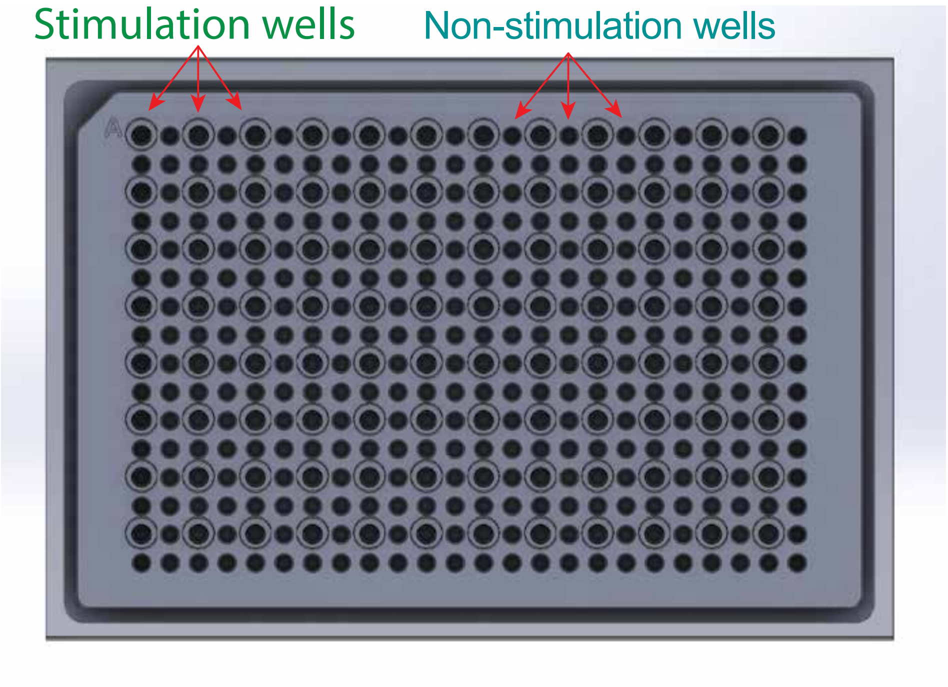


**B**

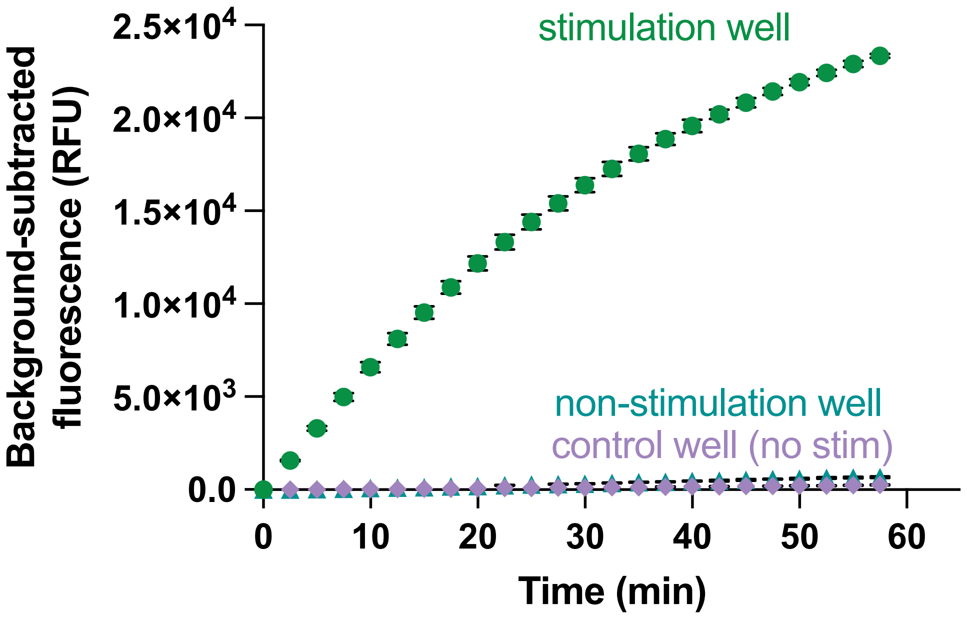


**Figure S5:** **No activity in non-stimulation wells. (A)** Top view of the 384 well plate with the acrylic custom plate adapter. **(B)** CRISPR reactions with a target concentration (1e6 cp/μL) were mixed simultaneously. Reactions in no stimulation wells were compared with reactions kept outside of the LUCas stimulation set up (control wells). Data points represent the mean ± SEM (n=3).

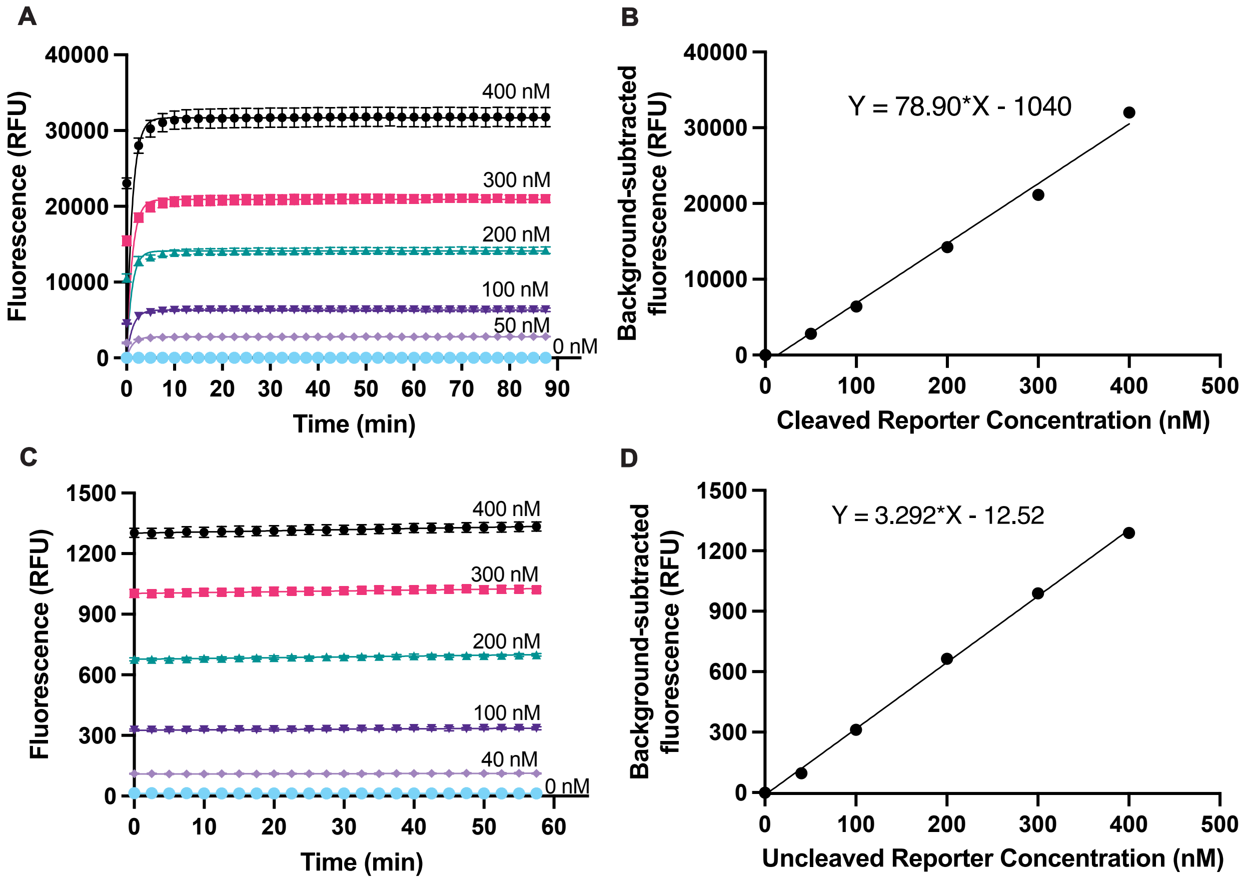


**Figure S6:** **Fluorescence versus cleaved and uncleaved reporter concentration calibration. (A)** Raw measurements of fluorescence over time for LbuCas13a with gRNA 1 and 1e7 cp/μL of RNA target concentration. **(B)** Background-subtracted fluorescence versus cleaved reporter concentration with a linear regression fit. **(C)** Raw measurements of fluorescence over time for uncleaved reporter concentrations. **(D)** Background-subtracted fluorescence versus uncleaved reporter concentration with a linear regression fit. In all panels, the data represents the mean ± SEM across three replicates.

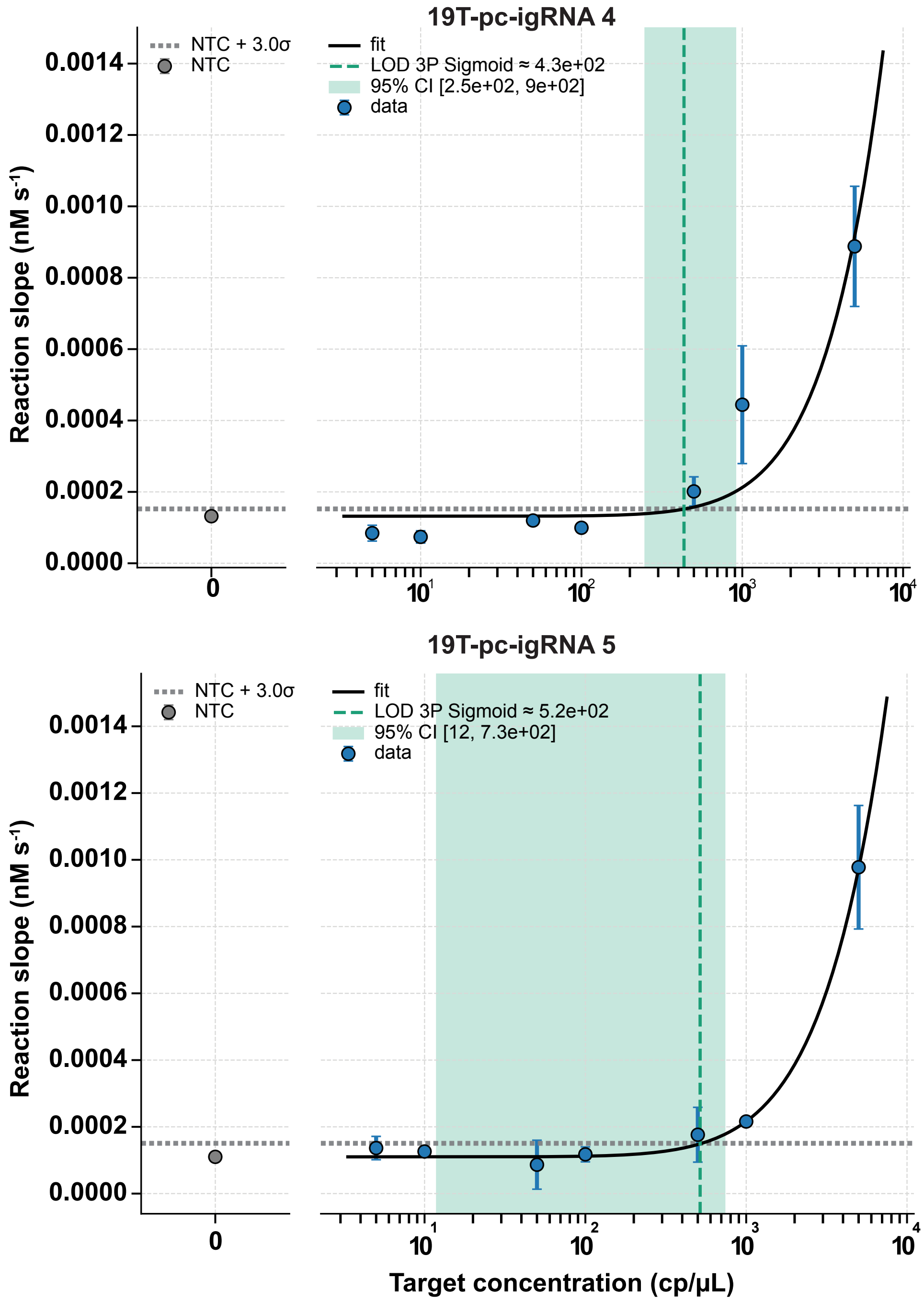


**Figure S7:** Limit-of-detection (LOD) analysis for 19T-pc-igRNA 4 and 19T-pc-igRNA 5 showing the measured background (NTC) and measured LOD in target concentration (vertical green dashed line) and 95% CI (green shaded region).

### Supplementary Tables

CRISPR Guides:

| Name | Sequence | Vendor |
|---|---|---|
| gRNA 1 | GACCACCCCAAAAAUGAAGGGGACUAAAAC**UUUGCGGCCAAUGUUUGUAA** | Synthego |
| gRNA 2 | GACCACCCCAAAAAUGAAGGGGACUAAAAC**GGUCCACCAAACGUAAUGCG** | Synthego |
| gRNA 3 | GACCACCCCAAAAAUGAAGGGGACUAAAAC**CAGUUGUGAUGAUUCCUAAG** | Synthego |
| gRNA 4 | GACCACCCCAAAAAUGAAGGGGACUAAAAC**AAGGUGUGACUUCCAUGCCA** | IDT |
| gRNA 5 | GACCACCCCAAAAAUGAAGGGGACUAAAAC**AUUGGUGUUAAUUGGAACGC** | IDT |
| gRNA FluA | GACCACCCCAAAAAUGAAGGGGACUAAAAC**CAGCUGACAUGAGUAUUGGA** | IDT |

Photocleavable Guides:

| Name | Sequence | Vendor |
|---|---|---|
| 19T-pc-igRNA 1 | TTTTTTTTTTTTTTTTTTT/iSpPC/CGACCACCCCAAAAAUGAAGGGGACUAAAAC**UUUGCGGCCAAUGUUUGUAA** | IDT |
| 9T-pc-pigRNA 1 | TTTTTTTTT/iSpPC/CGACCACCCCAAAAAUGAAGGGGACUAAAAC**UUUGCGGCCAAUGUUUGUAA** | IDT |
| 9T-pc-pigRNA 2 | TTTTTTTTT/iSpPC/CGACCACCCCAAAAAUGAAGGGGACUAAAAC**GGUCCACCAAACGUAAUGCG** | IDT |
| 9T-pc-pigRNA 3 | TTTTTTTTT/iSpPC/CGACCACCCCAAAAAUGAAGGGGACUAAAAC**CAGUUGUGAUGAUUCCUAAG** | IDT |
| 19T-pc-igRNA 4 | TTTTTTTTTTTTTTTTTTT/iSpPC/CGACCACCCCAAAAAUGAAGGGGACUAAAAC**GGUCCACCAAACGUAAUGCG** | IDT |
| 19T-pc-igRNA 5 | TTTTTTTTTTTTTTTTTTT/iSpPC/CGACCACCCCAAAAAUGAAGGGGACUAAAAC**CAGUUGUGAUGAUUCCUAAG** | IDT |

Targets and Reporter:

| Name | Sequence | Vendor |
|---|---|---|
| gRNA 1 target | GGAACUGA**UUACAAACAUUGGCCGCAAA**UUGCACAA | Synthego |
| gRNA 2 target | UGCACCC**CGCAUUACGUUUGGUGGACC**CUCAGAUU | Synthego |
| gRNA 3 target | CUUGUUUU**CUUAGGAAUCAUCACAACUG**UAGCUGCA | Synthego |
| gRNA 4 target | TCGCGCAT**TGGCATGGAAGTCACACCTT**CGGGAACG | Synthego |
| gRNA 5 target | AGGACAAG**GCGTTCCAATTAACACCAAT**AGCAGTCC | Synthego |
| PolyU Reporter | /56-FAM/UUUUU/3IABkFQ/ | IDT |

DNA is underlined. The 20-nucleotide sequence in bold is complementary to the gRNA spacer

Table S1: **Oligonucleotides used in this study.**
